# Discover Your Potential: The Influence of Kinematics on a Muscle’s Ability to Contribute to the Sit-to-Stand Transfer

**DOI:** 10.1101/2021.02.26.433097

**Authors:** Sarah A. Roelker, Laura C. Schmitt, Ajit M.W. Chaudhari, Robert A. Siston

## Abstract

Existing methods for estimating how individual muscles contribute to a movement require extensive time and experimental resources. In this study we developed an efficient method for determining how changes to lower extremity joint kinematics affect the potential of individual muscles to contribute to whole-body center-of-mass vertical (support) and anteroposterior (progression) accelerations. A 4-link 2-dimensional model was used to assess the effect of kinematic changes on muscle function. Joint kinematics were systematically varied throughout ranges observed during the momentum transfer phase of the sit-to-stand transfer. Each muscle’s potential to contribute to support and progression was computed and compared to simulated potentials estimated by traditional dynamic simulation methods for young adults and individuals with knee osteoarthritis (KOA). The new method required 4-10s to compute muscle potentials per kinematic state and computed potentials were consistent with simulated potentials. The new method identified differences in muscle potentials between groups due to kinematic differences, particularly decreased anterior pelvic tilt in young adults, and revealed kinematic and muscle strengthening modifications to increase support. The methods presented provide an efficient, systematic approach to evaluate how joint kinematic adjustments alter a muscle’s ability to contribute to movement and can identify potential sources of pathologic movement and rehabilitation strategies.

## Introduction

A muscle induced acceleration analysis (IAA) identifies the contribution of an individual muscle to the acceleration of a body segment or a joint connecting two body segments [1]. An IAA has been used to determine how individual muscles facilitate movement through contributions to the vertical support and forward progression of the body’s center of mass (COM) during walking (e.g., Anderson and Pandy, 2003; Liu et al., 2006; Neptune et al., 2001), running (e.g., Hamner et al., 2010), the sit-to-stand (STS) transfer (e.g., Caruthers et al., 2016), and climbing stairs (e.g., Caruthers et al., 2018; Lin et al., 2015). In general, these studies determined that the gluteus maximus, quadriceps, and plantarflexors are the primary contributors to the acceleration of the COM during these tasks. However, individual muscle contributions differ between tasks due to differences in kinematics [1]. For example, during walking [4] the gluteus maximus contributes to slowing the forward progression of the COM during early stance, but contributes to forward propulsion when rising from a chair [6] and ascending stairs [7].

Determining how task demands as well as changes to the skeletal, muscular, and nervous systems due to aging, disease, or injury affect movement mechanics and muscle function is a key step toward improved care for individuals with mobility limitations. In individuals with knee osteoarthritis (KOA) the quadriceps are commonly weak [9], and, therefore, this muscle group is most frequently targeted for strengthening; doing so has been shown to reduce pain and improve function [10]. However, current KOA rehabilitation programs often leave a large number of individuals, up to 40% in some cases, without significant improvement in short-term pain or ability to perform daily activities [11]. Although muscle strength is an important component that contributes to a muscle’s ability to facilitate movement, the mechanisms that govern force production are functionally complex, and muscle strengthening may not be sufficient to improve an individual’s ability to perform activities of daily living, such as rising from a chair. For example, individuals with KOA exhibit altered kinematics during the STS transfer, including increased trunk flexion and decreased knee joint excursion [12]. These changes in joint kinematics will alter the force and torque generation capacity of the muscles that cross the joint by altering the muscle’s fiber length and moment arm, respectively. Indeed, a recent simulation study investigated what degree of simulated muscle weakness could be tolerated while maintaining the subject’s experimentally observed joint mechanics during the STS transfer and found that simulations of individuals who performed the STS transfer with greater knee flexion were less tolerant to quadriceps weakness than individuals who used decreased knee flexion to complete the task [13]. Thus, there is a relationship between muscle strength and the range of kinematics that enable an individual to use that strength to accomplish a movement task. To develop targeted therapies that enable patients to take advantage of their strength gains from rehabilitation and ultimately improve patient function, there is a need to understand the complex interactions between kinematics, muscle strength, and muscle function.

Although an IAA approach can investigate the influence of altered kinematics, muscle forces, and inertial properties and their complex interactions on the ability of individual muscles to contribute to movement and ultimately mobility, unfortunately, the current implementation of IAA has two significant shortcomings that limit our ability to take this key step in patient care. Existing methods for determining muscle induced accelerations require extensive 1) experimental resources and 2) computational time, hindering the application of important insights from IAA in a clinical setting. A more straightforward and time efficient method of determining how modifications to kinematics and muscle strength will impact motion could improve the identification of appropriate rehabilitation targets to increase a patient’s mobility. Moreover, IAA is currently used as a post-hoc analysis of an observed motion and thus, in its current form, cannot be readily used as a predictive tool to guide treatment based on the complex changes to both muscle strength and movement patterns as a result of rehabilitation. If the current IAA approach could be adapted to develop a faster method that requires minimal resources and systematically evaluates how changes to an individual’s kinematics would affect the ability of a muscle to contribute to the movement, it would be possible to predict required strength changes to achieve a new movement pattern or, conversely, predict kinematic modifications that would leverage an individual’s current strength. Ultimately, such an approach could identify, based on a patient’s current strength levels and movement limitations, the combination of kinematic and strength modifications to be targeted during rehabilitation to optimize a particular musculoskeletal parameter (e.g., minimizing muscle activation demand).

Therefore, the purpose of this study was to develop a more efficient IAA method to determine how changes to lower extremity joint kinematics affect the ability (potential) of individual muscles to contribute to COM acceleration. The practical application of the new method was demonstrated in an analysis of muscle potentials for kinematic states observed during the momentum transfer phase of the STS transfer [14]. The momentum transfer phase was chosen because the greatest net muscle contributions to support are required in this phase [6]. The muscle potentials computed by this new method were compared to those estimated by traditional musculoskeletal modeling and simulation methods for unimpaired young adults (YA) and individuals with end-stage KOA during the momentum transfer phase of the STS transfer. Finally, the sensitivity of muscle potentials to common kinematic modifications during the STS including changes in foot position [15], pelvis height (as a surrogate for seat height [16]), lumbar flexion [17], and pelvic tilt [18] are presented.

## Materials & Methods

### Model

A four link 2-dimensional sagittal plane model (2D model; Figure 1A) with four revolute joints was used to assess the effects of various STS kinematic states on the function of 23 lower extremity muscles. The foot was modeled as a triangle, such that the three sides of the triangle are defined by vectors between three vertices located at the posterior calcaneus (C), the distal toe (T), and the ankle joint (A). The origin, O, of the coordinate reference frame was defined as the intersection point of the base of the foot, CT, and a perpendicular vector from CT to A. The origin was set coincident with the ground such that the positive X axis was directed anteriorly, the positive Y axis was directed superiorly, and positive rotations were defined in the counterclockwise direction about the positive Z axis (directed out of the page). Link 0 was formed by a rigid, perpendicular connection between CT, and a vector from O to A. Link 1, defined as the vector from A to the knee joint (K), represents the shank. Link 2, defined as the vector from K to the hip joint (H), represents the thigh. Link 3 represents a lumped model of the head, arms, and torso (HAT) and was defined as the vector from H to the HAT center of mass. The four revolute joints at points O, A, K, and H were parameterized by four generalized coordinates *θ*_0_, *θ*_1_, *θ*_2_, and *θ*_3_, respectively. Coordinate *θ*_0_ was defined as the angle between the X axis and CT. Given a 90° counterclockwise rotation from CT to OA, *θ*_1_ was defined as the angle between OA and Link 1. Finally, *θ*_2_ and *θ*_3_ were defined as the angles between Links 1 and 2 and between Links 2 and 3, respectively.

**Figure 1:**
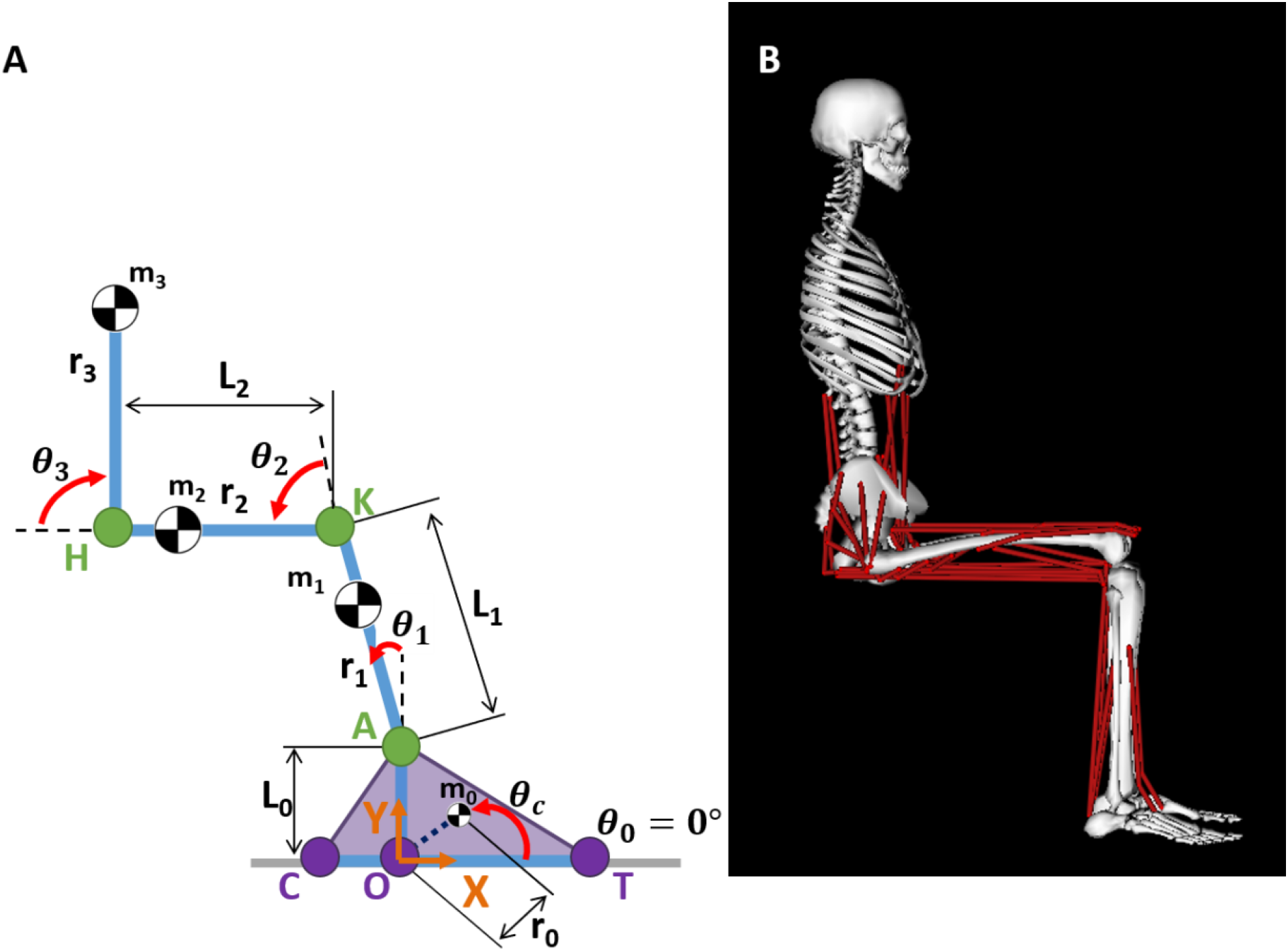
A) Four-link 2D model. Reference frame defined with origin at O, with the positive X axis directed anteriorly coincident with the ground (grey line), the positive Y axis directed superiorly, and positive rotation defined in the counterclockwise direction. Link 0 is formed by a rigid, perpendicular connection between lines CT and OA. Link 1 is defined by line AK. Link 2 is defined by line KH. Link 3 is a lumped representation of the head, arms, and torso (HAT) defined by the line from H to the center of mass of the HAT. Generalized coordinates: *θ*_0_, *θ*_1_, *θ*_2_, and *θ*_3_. Variables L_i_, r_i_, and mi define the lengths, center of mass location relative to the proximal joint, and mass, respectively, of segment *i*. *θ*_*c*_ defines the angle between CT and the vector from O to the center of mass of Link 0. B) Musculoskeletal (MSK) model.

The masses, lengths, center of mass (COM) locations, and moments of inertia of the segments were defined based on a generic 12 segment, 23 degree of freedom musculoskeletal model (MSK; Figure 1B) [19], modified to include a reduced muscle set (46 musculotendon actuators). The distances, ri, between the joint center and the COM of the segments, and the segment lengths, Li, were determined from the MSK model. The masses of the torso and pelvis from the MSK model were lumped together in the HAT segment of the 2D model. Similarly, the mass of the foot in the 2D model was calculated as the sum of the masses of the talus, calcaneus, and toes in the MSK model. The COM of the HAT and the COM of the foot were calculated by dividing the sum of the product of the individual masses of each body and their COM locations by the total mass. The moments of inertia of the HAT and the foot were determined using the parallel axis theorem. Muscle lengths and moment arms were determined for each muscle using the muscle properties defined in the MSK model.

### Calculation of Individual Muscle Potential to Contribute to COM Acceleration

The equations of motion of the model are defined by

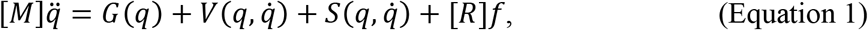

where *M* represents the mass matrix, *q* represents the generalized coordinates, *G* represents the gravitational force, *V* represents the force due to Coriolis and centrifugal effects, *S* represents the intersegmental forces, and *f* is an array containing the muscle forces transformed to generalized forces by matrix *R*, containing the muscle moment arms. The induced acceleration of any force acting on the system is

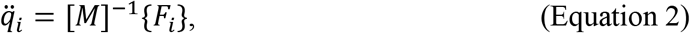

where *F*_*i*_ is the contribution of any force (gravitation, Coriolis, contact, or muscle) to the acceleration. To determine the acceleration solely due to the application of a muscle force (muscle induced acceleration), *G*(*q*), 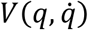, and 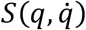 can be dropped from the equation such that Equation 1 can be simplified to

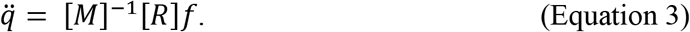

In Equation 3, [*M*] is an *n* × *n* matrix, [*R*] is an *n* × *m* matrix and [*f*] is an *m* × 1 matrix, where *n* is the number of generalized coordinates in the model and *m* is the number of muscles in the model.

A muscle’s potential to contribute to the support and progression of the body COM represents a muscle’s contribution to acceleration per Newton of force. In this study, individual muscle potentials were calculated as

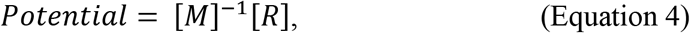

where [*M*] was a 4 × 4 matrix (*n* = 4 generalized coordinates) and [*R*] was a 4 × 23 matrix (*m* = 23 muscles).

### Mass Matrix Derivation

The mass matrix of the model was derived for the 2D model with four revolute joints using the Lagrange formulation for defining the equations of motion [20]. The generic form of [*M*] was obtained by

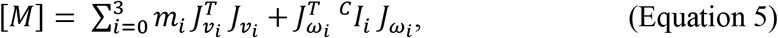

where *m*_*i*_ is the mass of link *i*, ^*c*^*I*_*i*_ is the inertial tensor of link *i*, 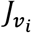 is the Jacobian matrix corresponding to the linear motion of the COM of link *i*, 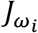 is the Jacobian matrix corresponding to the angular motion of link *i*, and 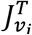 and 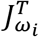 are the transpose of 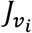 and 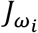, respectively.

Given the position vector of the COM of link 0 in frame {0} 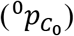

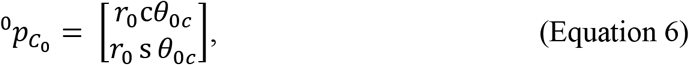

where *θ*_*c*_ is the angle between CT and the vector from O to the center of mass of link 0 (Figure 1A), c*θ*_0*c*_ = cos (*θ*_0_ + *θ*_*c*_), and s*θ*_0*c*_ = sin(*θ*_0_ + *θ*_*c*_), the Jacobian matrix corresponding to the linear motion of the COM of link 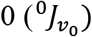 is obtained by direct differentiation of 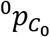 :

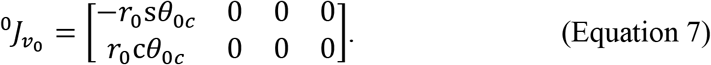

Similarly, the Jacobian matrix corresponding to the linear motion of the COM of link 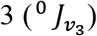 was obtained by direct differentiation of the position vector of the COM of link 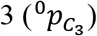

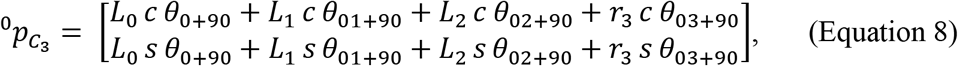

where c*θ*_03+90_ = cos(*θ*_0_ + *θ*_1_ + *θ*_2_ + *θ*_3_ + 90) and s *θ*_03+90_ = sin(*θ*_0_ + *θ*_1_ + *θ*_2_ + *θ*_3_ + 90), which gives, in frame {0},

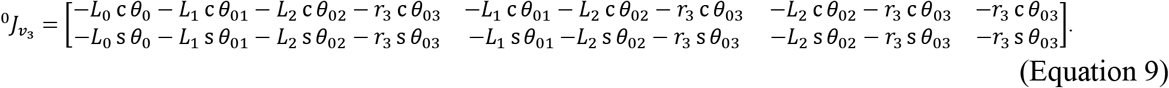

The Jacobian matrix corresponding to the angular motion of the COM of link 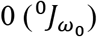 was obtained by

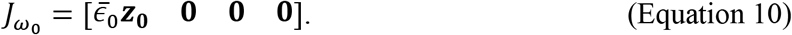

In frame {0}, this matrix is

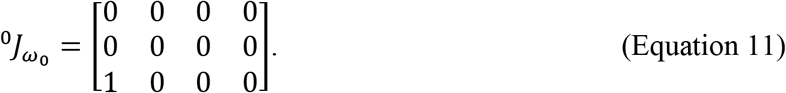

Similarly, the Jacobian matrix corresponding to the angular motion of the COM of link 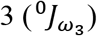 was obtained by

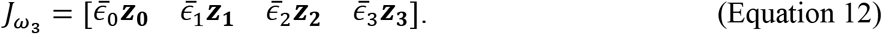

In frame {0}, this matrix is

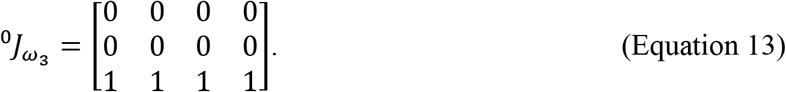

### Kinematic States

Kinematics were varied at each joint throughout ranges that are commonly observed in the momentum transfer phase of the STS transfer, which is defined as the phase that begins the moment the individual is no longer in contact with the seat and ends at the time of maximum ankle dorsiflexion [14]. To compute [*R*] for each kinematic position, moment arms were determined from the MSK model in OpenSim by converting the joint positions of the kinematic state from the 2D model joint angle definition to the MSK model joint angle definition using the following equations:

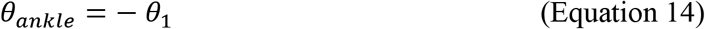

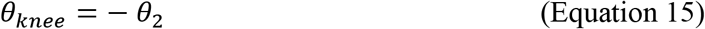

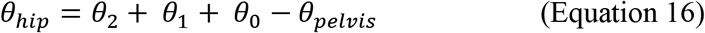

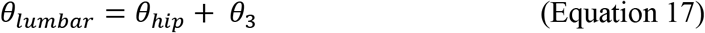

where *θ*_*ankle*_, *θ*_*knee*_, *θ*_*hip*_, *θ*_*lumbar*_, and *θ*_*pelvis*_ are the positive joint positions (ankle dorsiflexion, knee extension, hip flexion, lumbar extension, and posterior pelvic tilt, respectively) in the MSK model. All other rotational degrees of freedom in the MSK model were prescribed to be 0°.

The initial kinematic position of the 2D model was *θ*_0_ = 0°, *θ*_1_ = 0°, *θ*_2_ = 90°, and *θ*_3_ = −90°. Except for *θ*_0_, these angles and the pelvic tilt angle of the MSK model, were adjusted in 5° increments within specified ranges (Table 1) based on Schenkman, et al. (1990). The kinematic states were constrained such that the foot remained flat on the floor (*θ*_0_ = 0° for all trials) and the lumbar angle of the MSK model was restricted to between 90° of extension and 90° of flexion and the hip angle was limited to no more than 120° of flexion. Given the joint angle ranges and constraints, we calculated individual muscle potentials for support and progression for 14,758 unique kinematic states.

**Table 1:**
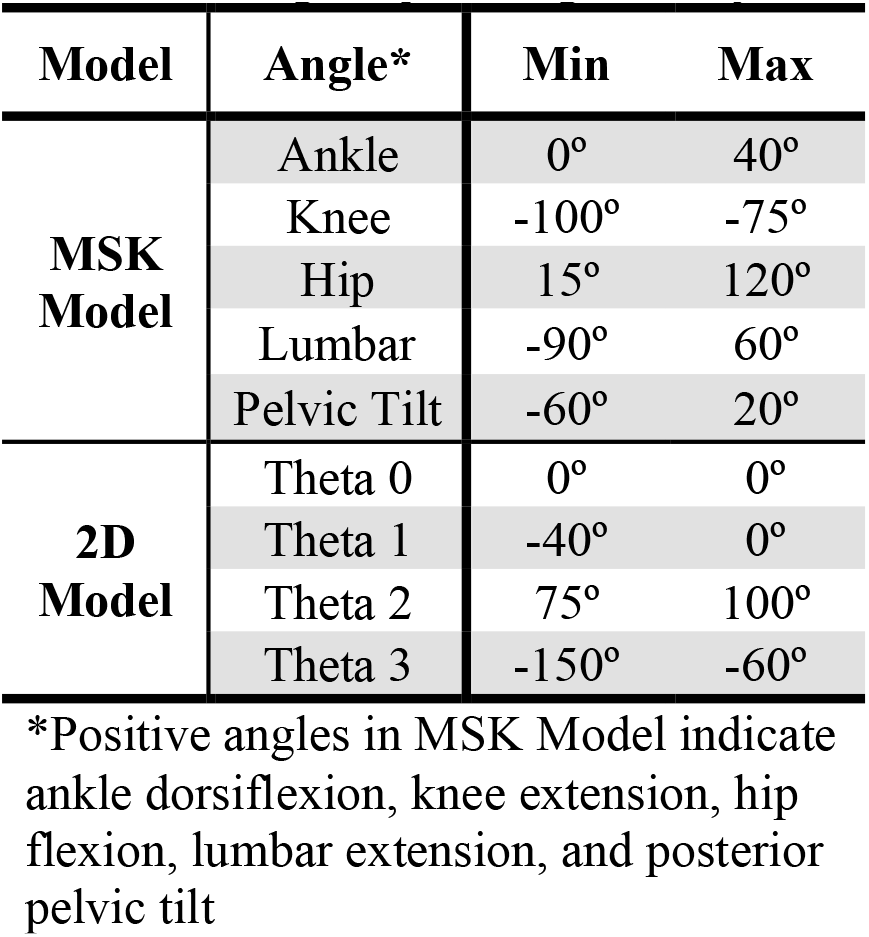
Range of joint angles analyzed

| Model | Angle* | Min | Max |
| --- | --- | --- | --- |
| **MSK Model** | Ankle | 0° | 40° |
|  | Knee | -100° | -75° |
|  | Hip | 15° | 120° |
|  | Lumbar | -90° | 60° |
|  | Pelvic Tilt | -60° | 20° |
| **2D Model** | Theta 0 | 0° | 0° |
|  | Theta 1 | -40° | 0° |
|  | Theta 2 | 75° | 100° |
|  | Theta 3 | -150° | -60° |

\*Positive angles in MSK Model indicate ankle dorsiflexion, knee extension, hip flexion, lumbar extension, and posterior pelvic tilt

### Transformation of Potentials from Joint Space to Task Space

To determine the individual muscle contributions to progression and support, defined along the X and Y Cartesian axes, respectively, the potentials were transformed from joint space to the Cartesian task space. The relationship between joint space (θ) and Cartesian space (X) can be defined using the definition of the Jacobian,

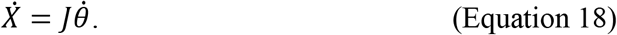

Differentiating Equation 18, the Cartesian acceleration is calculated as,

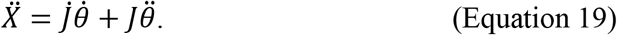

The potential of the muscles to contribute to joint accelerations were calculated under static conditions; therefore, the angular velocity of each joint angle is zero, giving,

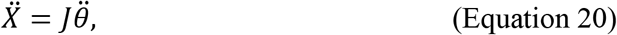

which was used to calculate the individual muscle potentials to contribute to progression and support for a given kinematic state.

### Muscle Potential Analysis

The time required to determine the muscle potentials for a single state was measured to estimate the computational time of the presented methods. Analyses were performed on a Lenovo Yoga C930 with an 8^th^ generation i7 Intel Core processor. Based on previous research of muscle function during the STS [6], the support and progression potentials of four primary contributors (gluteus maximus, soleus, vasti, and rectus femoris) and three secondary contributors (biceps femoris (long head), gastrocnemius (medial and lateral heads), and tibialis anterior) were chosen for further analysis. Muscle potentials were analyzed with respect to the position of the feet relative to the location of the hip, pelvis height, lumbar flexion angle, and pelvic tilt angle. Foot position (Figure 2) was calculated as the horizontal distance from the hip to the ankle:

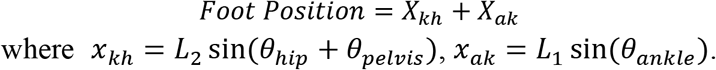

**Figure 2:**
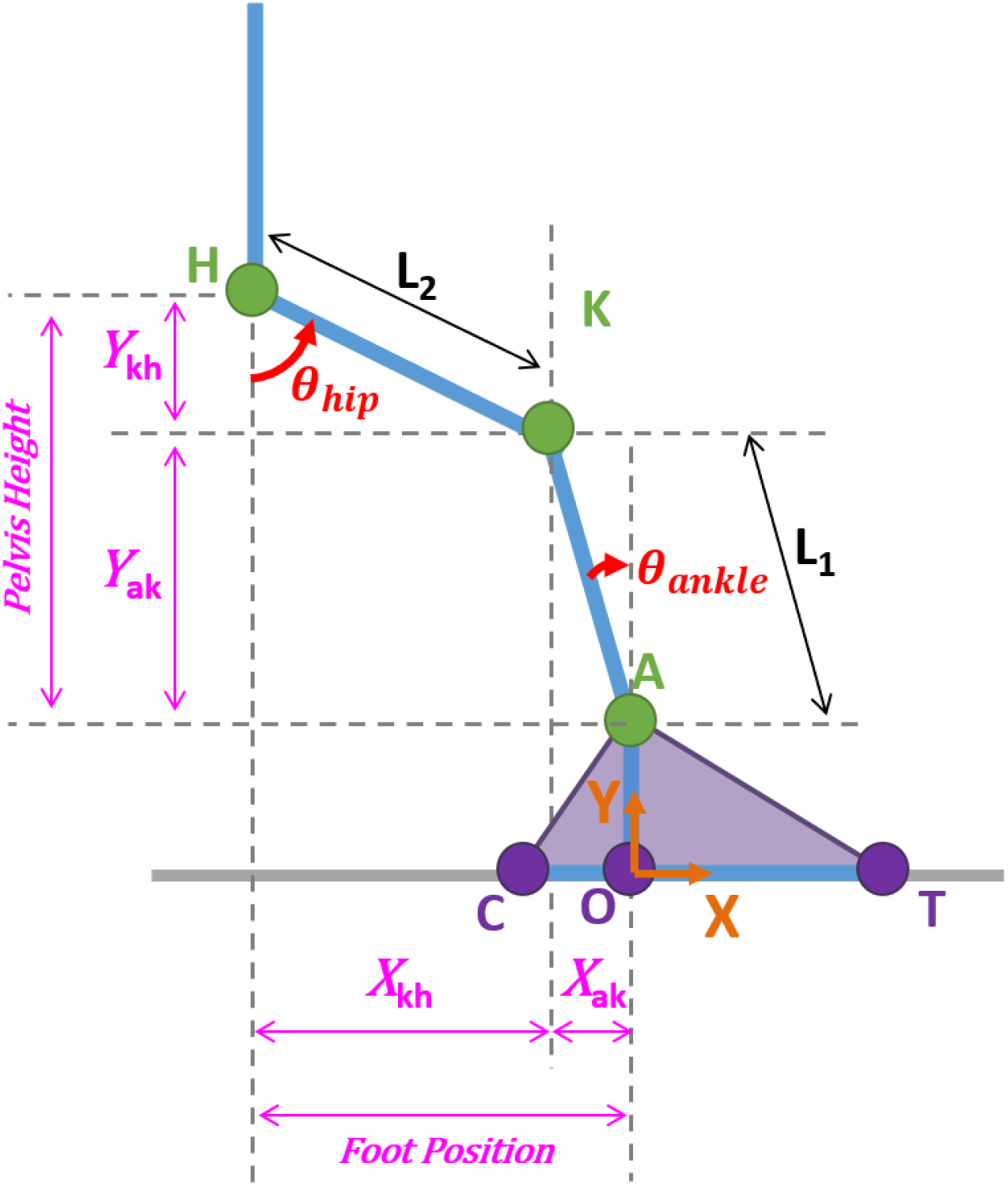
Dimensions for foot position and pelvis height calculations. X_kh_ and Y_kh_ are the horizontal and vertical distances, respectively, between K and H. X_ak_ and Y_ak_ are the horizontal and vertical distances, respectively, between A and K. Foot position is the sum of X_kh_ and X_ak_ and defined relative to H. Pelvis height is the sum of Y_kh_ and Y_ak_ and defined relative to A. Foot position and pelvis height are normalized to leg length (L_1_ + L_2_). In this figure, *θ*_*hip*_ is defined assuming a neutral pelvis tilt angle (*θ*_*pelvis*_ = 0°).

Foot position was normalized to leg length (L_1_ + L_2_) and is therefore a unitless metric. A foot position of 0 indicates the ankle is in line with the hip. The height of the pelvis relative to the ankle was defined (Figure 2) and calculated as the vertical distance from the ankle to the hip:

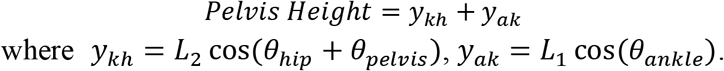

Pelvis height was also normalized to leg length. The muscle potentials computed by the presented (new) method were compared to potentials estimated using experimental 3D motion capture, modeling, and simulation (traditional) methods for seven YA (three female; 21.9 ± 2.3 years; 72.8 ± 11.4 kg; 1.74 ± 0.08 m) and nine individuals with KOA (six female; 60.6 ± 7.3 years; 84.9 ± 8.1 kg; 1.65 ± 0.10 m) based on the kinematics used by each cohort during the momentum transfer phase of the STS (Figure 3) [6,21]. For each muscle and each kinematic modification (foot position, pelvis height, lumbar flexion, and pelvis tilt), Pearson correlations assessed the strength of the relationship between a change in the kinematic modification and a change in the muscle’s support and progression potentials. The significance level, α was set to 0.05 *a priori*. Based on the correlation coefficient (*r*), correlations are categorized as very weak (*r* < 0.2), weak (0.20 ≤ r ≤ 0.39), moderate (0.40 ≤ r ≤ 0.59), strong (0.6 ≤ r ≤ 0.79) and very strong (0.8 ≤ r).

**Figure 3:**
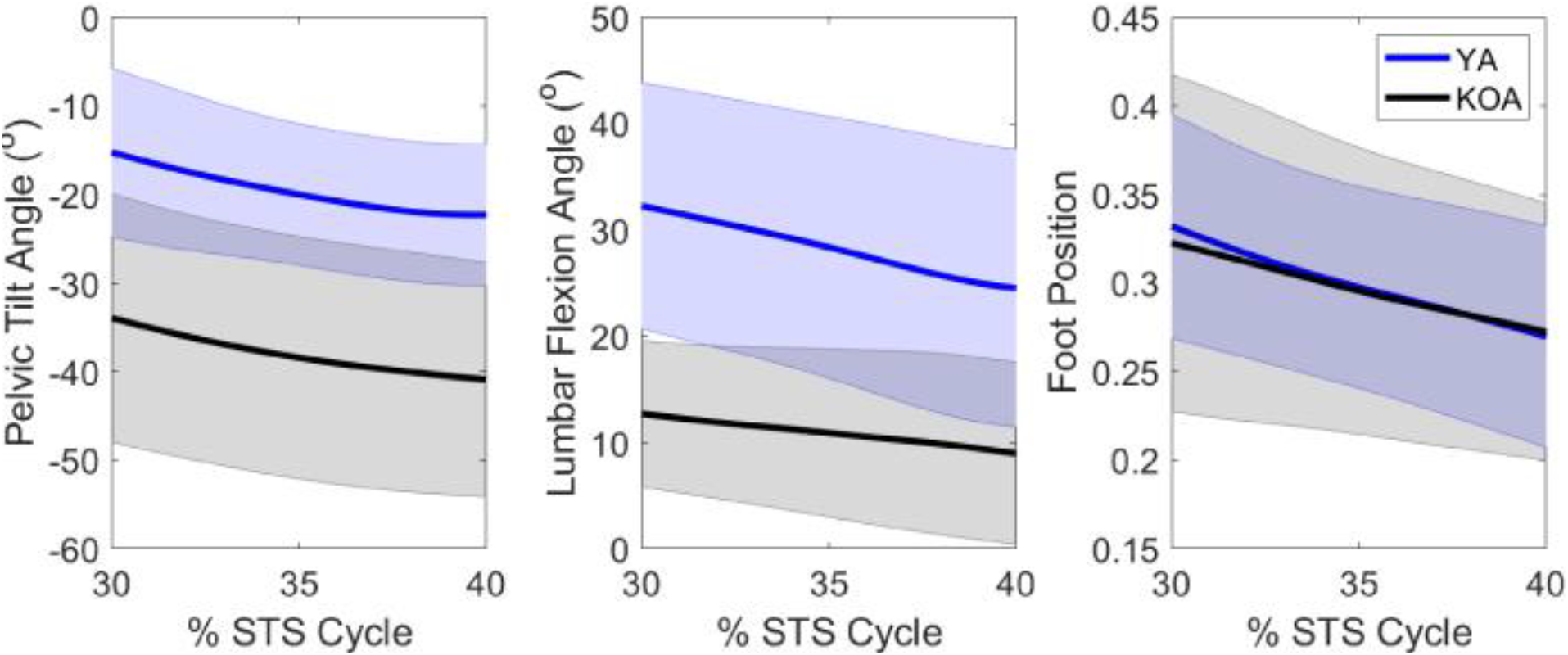
Experimental kinematics of YA and patients with KOA for key parameters during the momentum transfer phase of the STS transfer. More positive values indicate posterior pelvic tilt, lumbar flexion, and a more anterior foot position. The line represents the population mean and the shaded area represents ± one standard deviation.

## Results

### Efficiency and Verification of New Method

The new method calculated the support and progression potentials of 46 musculotendon actuators for a single kinematic state in 4-10s. Given the range of kinematic states exhibited by the YA and individuals with KOA during the momentum transfer phase, the support (Figure 4) and progression (Figure 5) potentials computed by the new method compared well with the muscle potentials estimated by the traditional methods. For each population, the range of magnitudes of the computed potentials typically fell within one standard deviation of the simulated potentials. Moreover, given the different ranges of pelvic tilt angles exhibited by each cohort, differences in computed muscle potentials were computed by the new method and verified by the simulated potentials. For example, for smaller anterior pelvic tilt angles the new method computed larger support potentials for the gluteus maximus and biceps femoris and smaller rectus femoris support potentials (Figure 4). The anterior pelvic tilt of the YA (−19.6° ± 8.1°) was less than the individuals with KOA (−38.0° ± 13.6°) and the YA had larger simulated support potentials for gluteus maximus (YA: 1.7 × 10^−3^ ± 0.7 × 10^−3^ m/Ns^2^; KOA: 0.7 × 10^−3^ ± 0.6 × 10^−3^ m/Ns^2^) and biceps femoris (YA: 0.3 × 10^−3^ ± 0.3 × 10^−3^ m/Ns^2^; KOA: −0.2 × 10^−3^ ± 0.3 × 10^−3^ m/Ns^2^) and smaller rectus femoris support potentials (YA: 0.3 × 10^−3^ ± 0.3 × 10^−3^ m/Ns^2^; KOA: 0.7 × 10^−3^ ± 0.3 × 10^−3^ m/Ns^2^). Similarly, the larger gluteus maximus progression potential of the YA compared to individuals with KOA (YA: 0.5 × 10^−3^ ± 0.3 × 10^−3^ m/Ns^2^, KOA: 0.2 × 10^−3^ ± 0.2 × 10^−3^ m/Ns^2^) was computed by the new method based on the smaller anterior pelvic tilt of the YA (Figure 5).

**Figure 4:**
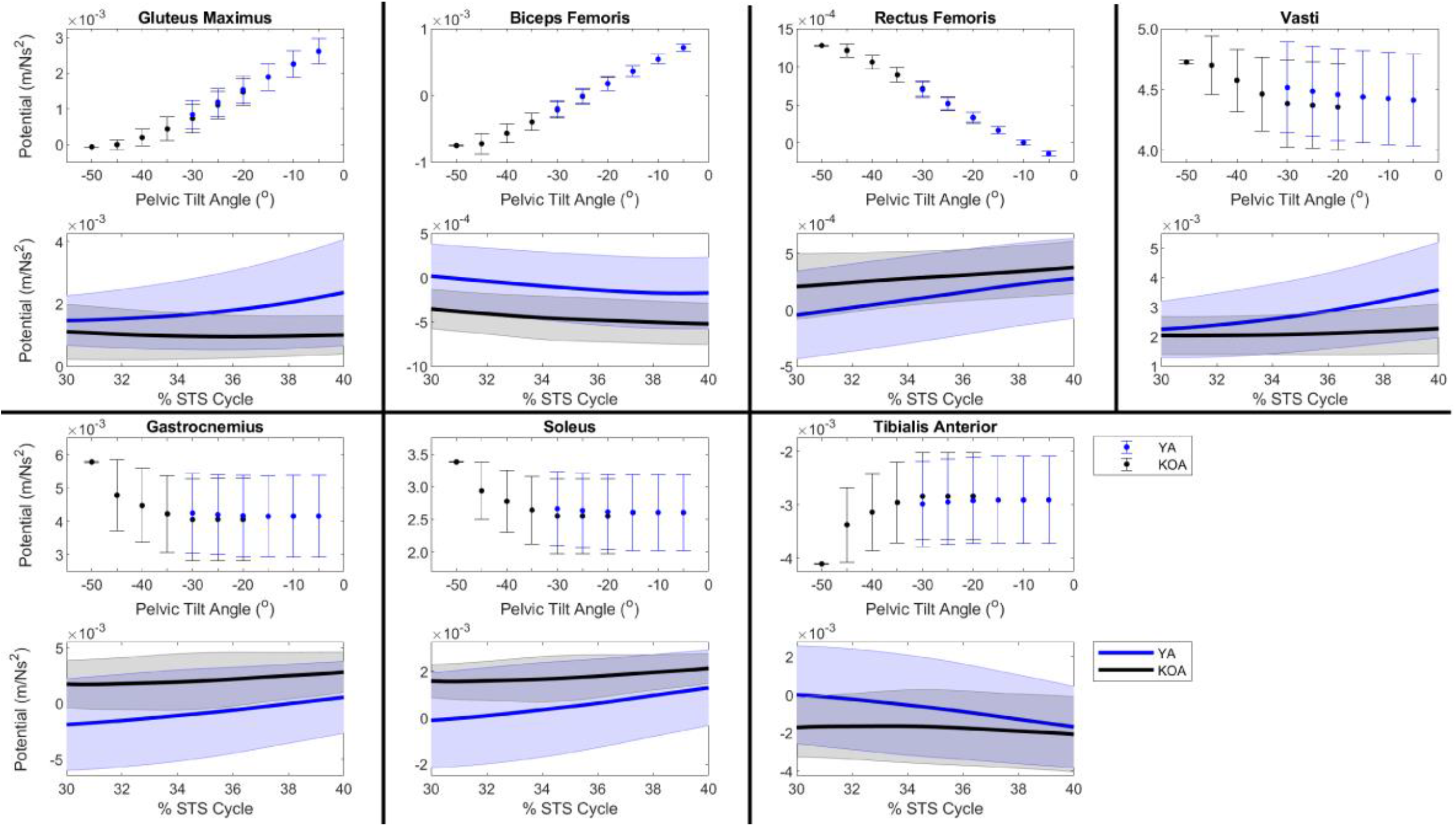
Muscle support potentials during the momentum transfer phase of the STS. For each muscle, the top plot expresses support potential as a function of pelvic tilt angle such that each data point represents the average muscle potential for a given pelvic tilt angle accounting for the range of kinematic states observed by the population with the respective pelvic tilt. Error bars represent ± one standard deviation from the average. Positive angles indicate posterior pelvic tilt. The bottom plot provides the time curve of the muscle’s support potential during the momentum transfer phase (30-40% of the STS cycle) estimated using a musculoskeletal model scaled to each subject’s anthropometrics in OpenSim. The line represents the population mean and the shaded area represents ± one standard deviation.

**Figure 5:**
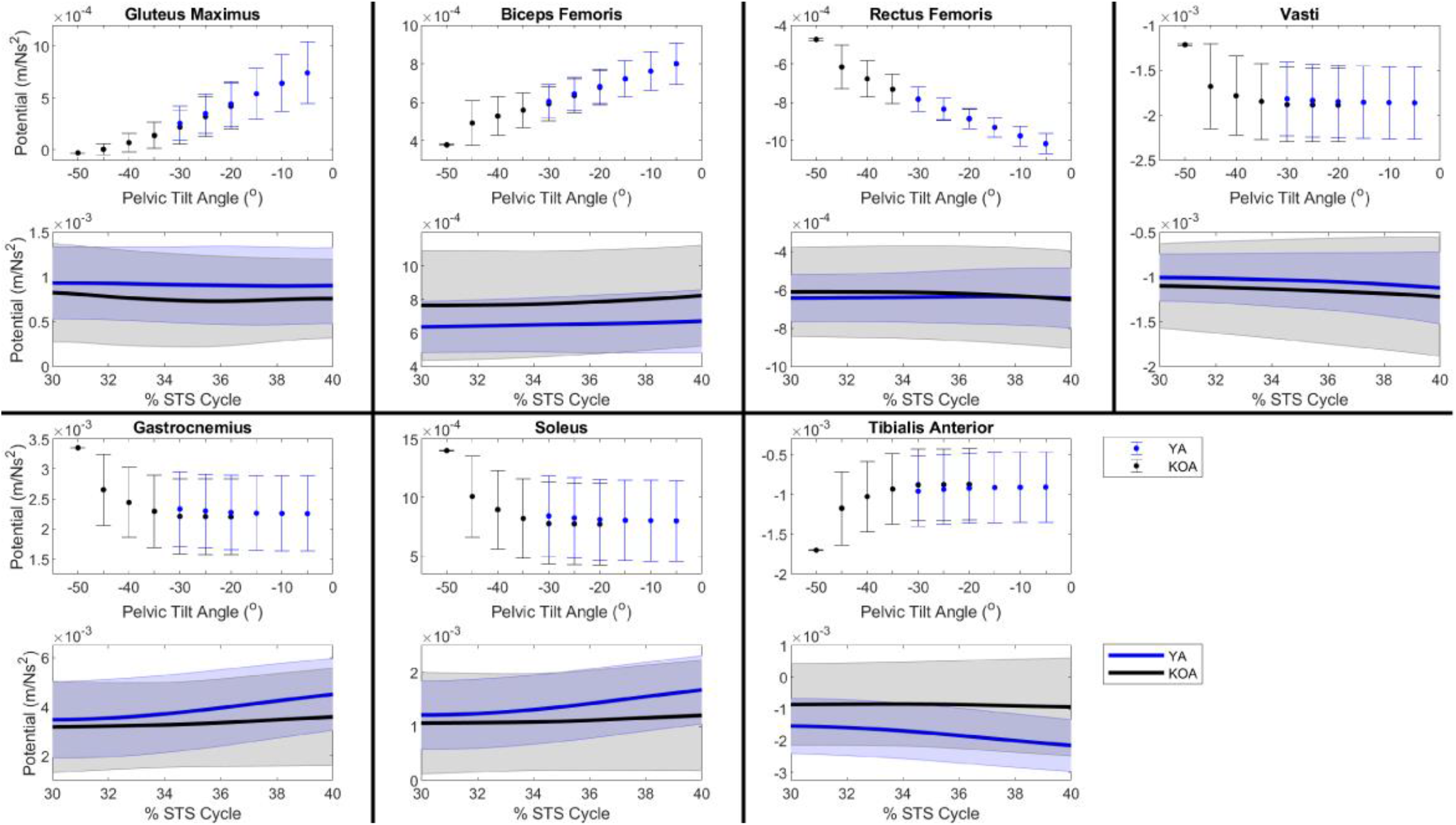
Muscle progression potentials during the momentum transfer phase of the STS. For each muscle, the top plot expresses progression potential as a function of pelvic tilt angle such that each data point represents the average muscle potential for a given pelvic tilt angle accounting for the range of kinematic states observed by the population with the respective pelvic tilt. Error bars represent ± one standard deviation from the average. Positive angles indicate posterior pelvic tilt. The bottom plot provides the time curve of the muscle’s progression potential during the momentum transfer phase (30-40% of the STS cycle) estimated using a musculoskeletal model scaled to each subject’s anthropometrics in OpenSim. The line represents the population mean and the shaded area represents ± one standard deviation.

### Sensitivity of Muscle Potentials to Kinematics

Support and progression potentials of all primary and secondary contributors are sensitive to changes in foot position, pelvis height, lumbar flexion angle and pelvic tilt angle to varying degrees as indicated by the significant correlations between all muscle potentials and kinematic modifications (Table 2). Pelvic tilt angle influences the lumbar flexion angle (Equations 16 and 17); therefore, the individual effects of foot position (Figure 6), pelvis height (Figure 7), and lumbar flexion angle (Figure 8) on muscle potentials are described for kinematics states with the pelvis in a neutral position and the effects of changes in pelvic tilt angle (Figure 9) are subsequently described considering all lumbar flexion angles and foot positions.

**Figure 6:**
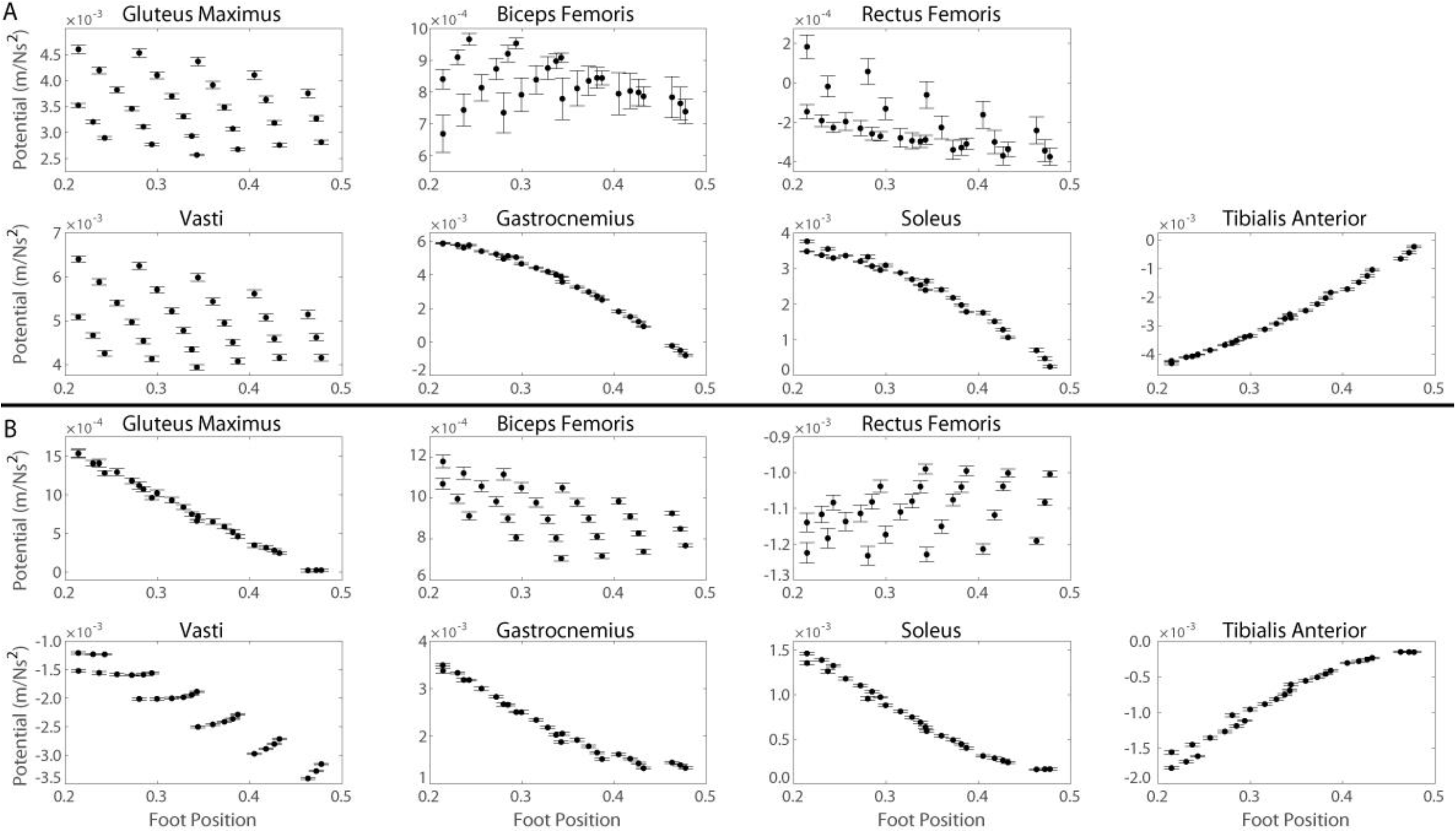
Muscle potentials to contribute to A) support and B) progression during the momentum transfer phase of the STS as a function of foot position. Each data point represents the average muscle potential for a given foot position accounting for the range of kinematic states with the respective foot position and a neutral pelvis position. Error bars represent ± one standard deviation from the average.

**Figure 7:**
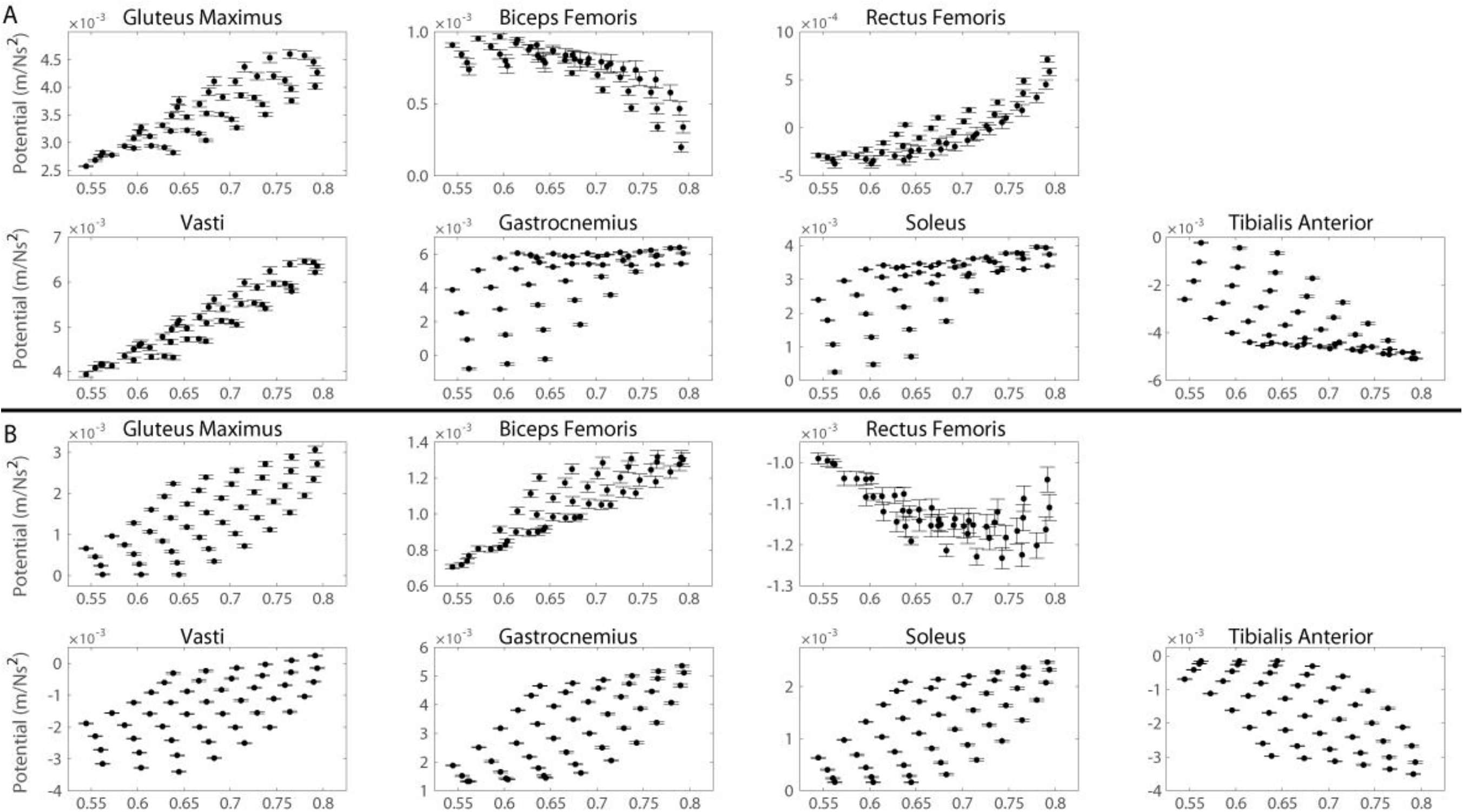
Muscle potentials to contribute to A) support and B) progression during the momentum transfer phase of the STS as a function of pelvis height. Each data point represents the average muscle potential for a given pelvis height accounting for the range of kinematic states with the respective pelvis height and a neutral pelvis position. Error bars represent ± one standard deviation from the average.

**Figure 8:**
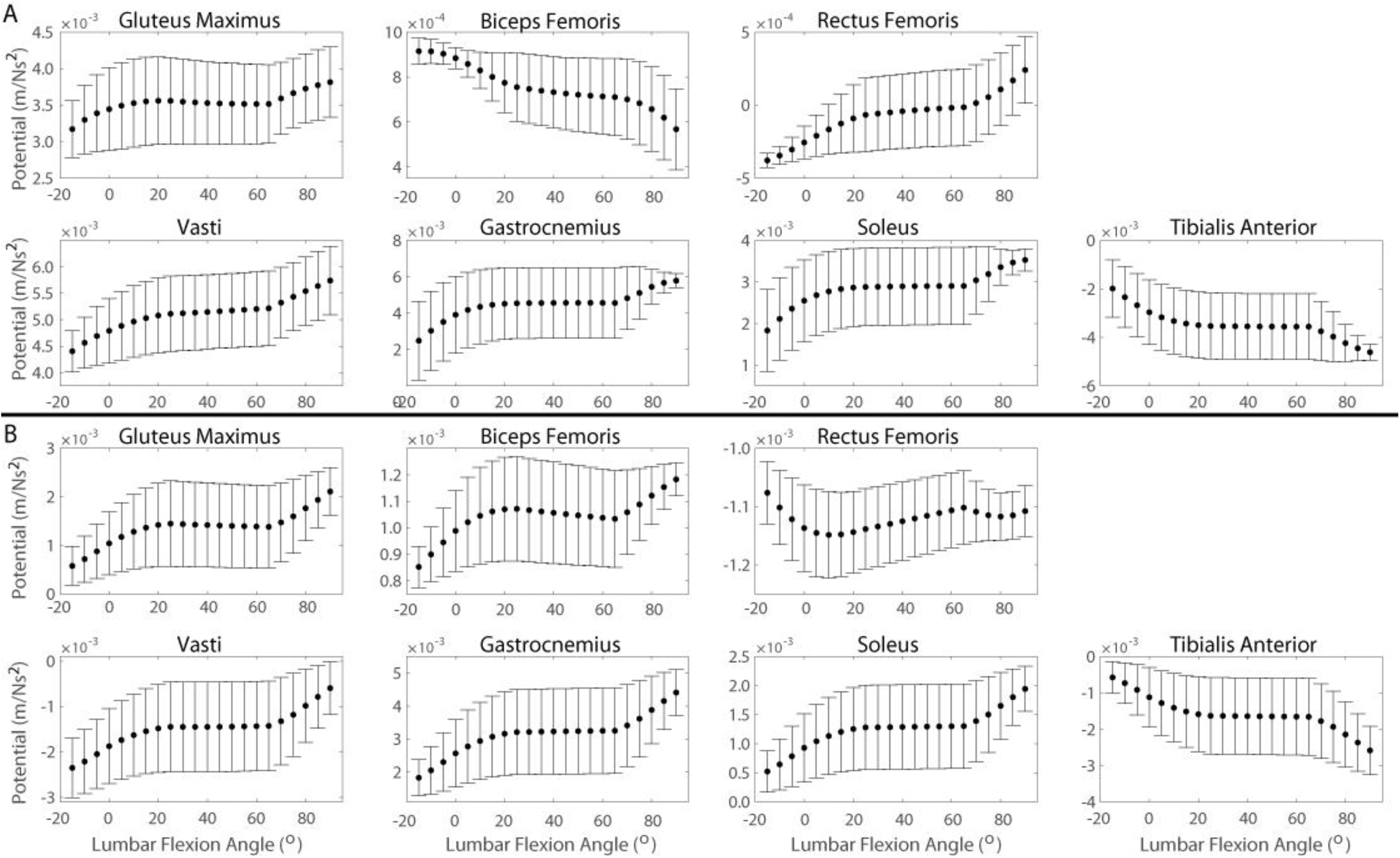
Muscle potentials to contribute to A) support and B) progression during the momentum transfer phase of the STS as a function of lumbar flexion angle. Each data point represents the average muscle potential for a given lumbar flexion angle accounting for the range of kinematic states with the respective lumbar angle and a neutral pelvis position. Error bars represent ± one standard deviation from the average. Positive angles indicate lumbar flexion.

**Figure 9:**
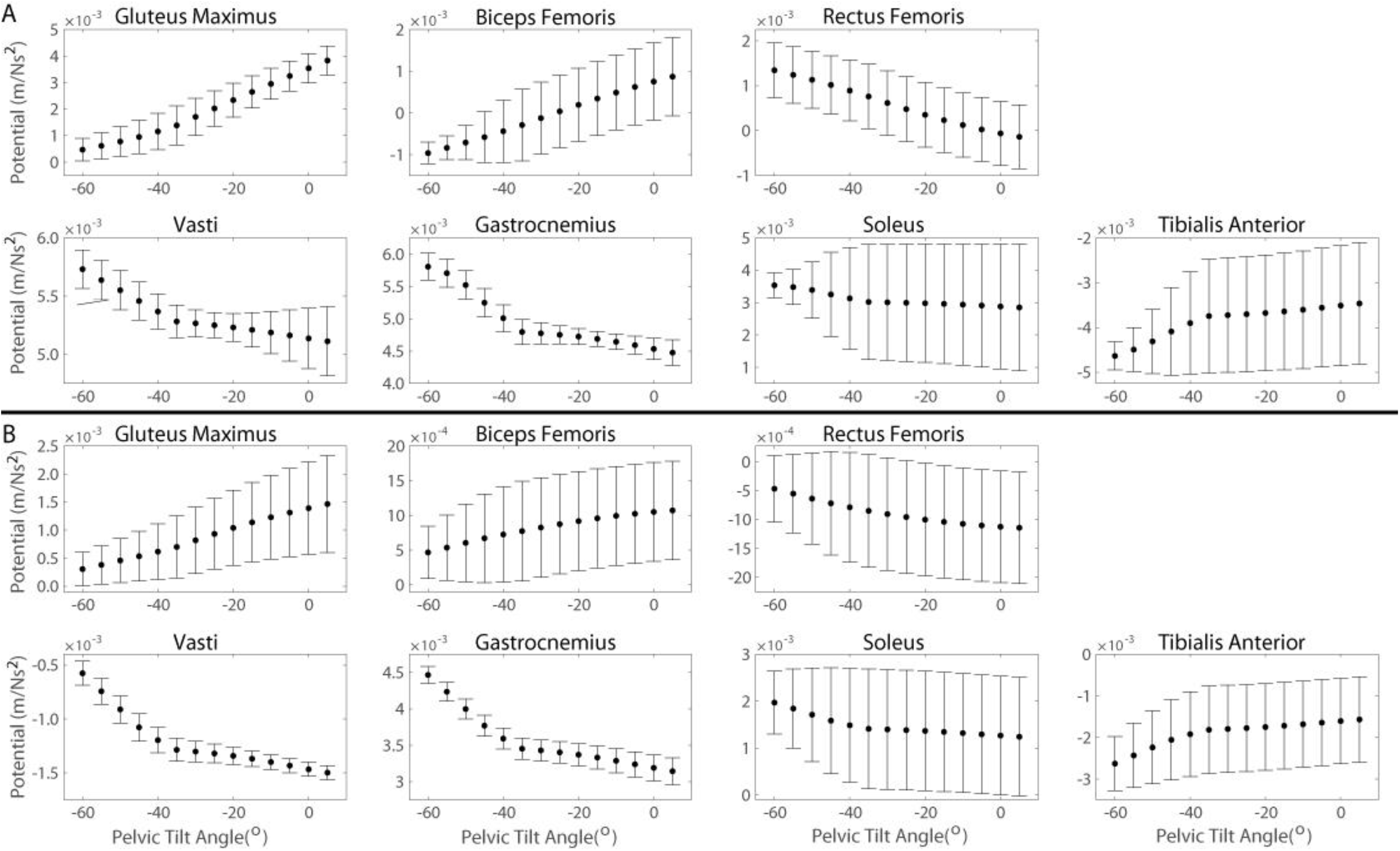
Muscle potentials to contribute to A) support and B) progression during the momentum transfer phase of the STS as a function of pelvic tilt angle. Each data point represents the average muscle potential for a given pelvic tilt angle accounting for the range of kinematic states. Error bars represent ± one standard deviation from the average. Positive angles indicate posterior pelvic tilt.

**Table 2:** Correlation coefficients for Pearson correlation between muscle potentials and kinematic modifications*

| Potential | Muscle | Foot Position | Pelvis Height | Lumbar Flexion | Pelvic Tilt |
| --- | --- | --- | --- | --- | --- |
| **Support** | Gluteus Maximus | -0.217 | 0.861 | 0.141 | 0.855 |
|  | Biceps Femoris | 0.612 | -0.758 | -0.443 | 0.953 |
|  | Rectus Femoris | -0.848 | 0.846 | 0.446 | -0.918 |
|  | Vasti | -0.394 | 0.939 | 0.328 | -0.195 |
|  | Gastrocnemius | -0.815 | 0.546 | 0.244 | -0.175 |
|  | Soleus | -0.800 | 0.640 | 0.268 | -0.189 |
|  | Tibialis Anterior | 0.916 | -0.648 | -0.290 | 0.206 |
| **Progression** | Gluteus Maximus | -0.998 | 0.684 | 0.259 | 0.460 |
|  | Biceps Femoris | -0.883 | 0.877 | 0.211 | 0.713 |
|  | Rectus Femoris | 0.280 | -0.643 | 0.127 | -0.871 |
|  | Vasti | -0.982 | 0.539 | 0.280 | -0.196 |
|  | Gastrocnemius | -0.990 | 0.662 | 0.319 | -0.219 |
|  | Soleus | -0.991 | 0.635 | 0.311 | -0.214 |
|  | Tibialis Anterior | 0.990 | -0.608 | -0.304 | 0.209 |

\* $p$ -value for all correlations: $p < 0.001$ . Positive (negative) correlation coefficients indicate muscle potentials increase (decrease) with more anterior foot placement, higher pelvis height, greater lumbar flexion, or a more posterior pelvic tilt.

For foot position, a positive (negative) correlation indicates that a more anterior (posterior) foot position is associated with increased muscle potential. Foot position had a very strong positive correlation with the tibialis anterior support potential and very strong negative correlations with the support potentials of rectus femoris, gastrocnemius, and soleus. Biceps femoris support potential had a strong positive correlation with foot position while there were weak correlations between foot position and the support potentials of gluteus maximus and vasti. The progression potentials of gluteus maximus, biceps femoris, vasti, gastrocnemius, and soleus had very strong negative correlations with foot position. The tibialis anterior progression potential had a very strong positive correlation with foot position. There was a weak positive correlation between foot position and the rectus femoris progression potential.

For pelvis height, a positive (negative) correlation indicates that a higher (shorter) pelvis height is associated with increased muscle potential. There were very strong positive correlations between pelvis height and the support potentials of gluteus maximus, rectus femoris, and gastrocnemius. The support potentials of biceps femoris and tibialis anterior have a strong negative correlation with pelvis height while the support potentials of soleus has a strong positive correlation with pelvis height. There was a moderate positive correlation between foot position and the gastrocnemius support potentials. The progression potentials of biceps femoris had a very strong positive correlation with pelvis height. Pelvis height had strong positive correlations with the progression potentials of gluteus maximus, gastrocnemius, and soleus and strong negative correlation with the progression potentials of rectus femoris and tibialis anterior. There was a moderate positive correlation between pelvis height and the vasti progression potentials.

For lumbar flexion angle, a positive (negative) correlations that a greater lumbar flexion (extension) angle is associated with increased muscle potential. Lumbar flexion angle had a moderate positive correlation with rectus femoris support potentials and a moderate negative correlation with biceps femoris support potentials. There were weak positive correlations between lumbar flexion angle and the support potentials of vasti, gastrocnemius, and soleus. There was a weak negative correlation between lumbar flexion angle and tibialis anterior support potentials and a very weak positive correlation between lumbar flexion angle and gluteus maximus support potentials. There were weak positive correlations between lumbar flexion angle and the progression potentials of gluteus maximus, biceps femoris, vasti, gastrocnemius, and soleus and a weak negative correlation between lumbar flexion angle and the progression potential of tibialis anterior. There was a very weak positive correlation between lumbar flexion angle and the rectus femoris progression potentials

For pelvic tilt angle, a positive (negative) correlation indicates that a more posterior (anterior) pelvic tilt angle is associated with increased muscle potential. There were very strong positive correlations between pelvic tilt angle and the support potentials of gluteus maximus and biceps femoris and a very strong negative correlation between pelvic tilt angle and rectus femoris support potentials. There was a weak positive correlation between tibialis anterior support potentials and pelvic tilt angle and very weak negative correlations between pelvic tilt angle and the support potentials of vasti, gastrocnemius, and soleus. A very strong negative correlation was observed between rectus femoris progression potentials and pelvic tilt angle while a strong positive correlation was observed between biceps femoris progression potentials and pelvic tilt angle. There were weak negative correlations between pelvic tilt angle and the progression potentials of gastrocnemius and soleus and a weak positive correlation between pelvic tilt angle and the tibialis anterior progression potentials. A very weak negative correlation was observed between pelvic tilt angle and vasti progression potentials.

## Discussion

The purpose of this study was to create an efficient yet robust method for establishing the relationship between lower extremity joint kinematics and individual muscle potentials to support and progression during activities of daily living and demonstrate its efficacy through an analysis of the sit to stand transfer. A computation speed of analyzing 5-15 kinematic states per minute was achieved. The typical frame rate of a video camera commonly used in a clinical setting to capture 2D joint angles is 60 frames (or states) per second and the average STS trial takes 1 second to complete. Thus, muscle potentials could be calculated for an entire trial in 12 minutes 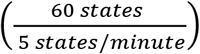 or less, which is considerably faster than traditional simulation techniques that require a few hours, rather than a few minutes, to obtain IAA results for a single trial. In addition, with the new IAA method, muscle potentials can be easily calculated for select movement phases, key events of a movement, or a range of feasible states that could be explored, which would reduce the number of states analyzed and further reduce the computational time.

We verified the potentials computed by the new method (computed potentials) against potentials estimated by traditional methods (simulated potentials). The computed muscle potentials were typically within one standard deviation of the simulated potentials (Figures 4 and 5). The 2D model’s segments’ masses and lengths were based on the generic MSK model and thus were not scaled to a specific individual. However, the mass (75.2 kg) and leg length (82.6 cm) of the generic MSK model falls within one standard deviation of the YA cohort’s average mass (72.8 ± 11.4 kg) and leg length (84.5 ± 3.3 cm) and the KOA cohort’s average leg length (81.1 ± 5.5 cm) and within two standard deviations of the KOA cohort’s average mass (84.9 ± 8.1 kg). The magnitudes of the simulated and computed muscle potentials are comparable, which suggests the new method is capable of reproducing simulated muscle potentials. Furthermore, despite the lack of subject specificity in the 2D model, the new method could accurately identify differences in muscle potentials between populations given the known differences in STS kinematics between YA and individuals with KOA.

The findings from this new method highlight the importance of the overall kinematic state of the body (quantified in the mass matrix) in determining how muscles facilitate movement. Modifying the body’s kinematic state, even when the position of the joint(s) crossed by a muscle remains the same, can alter a muscle’s potential. For example, with an anterior pelvic tilt of 15°, the gluteus maximus progression potential is 0.131 × 10^−3^ m/Ns^2^ (27.6%) larger at a foot position of 0.294 than a foot position of 0.337 with 5° greater knee and ankle flexion angles but the same hip flexion angle (Figure 10). Furthermore, although positioning the foot posteriorly has been shown to augment an individual’s ability to rise from a chair by reducing the peak hip extension moment (Janssen et al., 2002), our findings suggest that a considerable change in foot position may not necessarily affect a muscle’s potential to contribute to the movement. For example, given 5° of lumbar flexion and a neutral pelvis position, moving the foot to a 0.146 (40.6%) more posterior foot position from 0.360 to 0.214 has a negligible effect on the biceps femoris support potential (0.873 × 10^−3^ m/Ns^2^ and 0.885 × 10^−3^ m/Ns^2^, respectively) (Figure 10). In this case, the overall kinematic state remains relatively similar with the largest change in joint angles between foot positions occurring at the ankle (15°) compared to smaller changes at the joints crossed by the biceps femoris, the hip (5°) and knee (10°).

**Figure 10:**
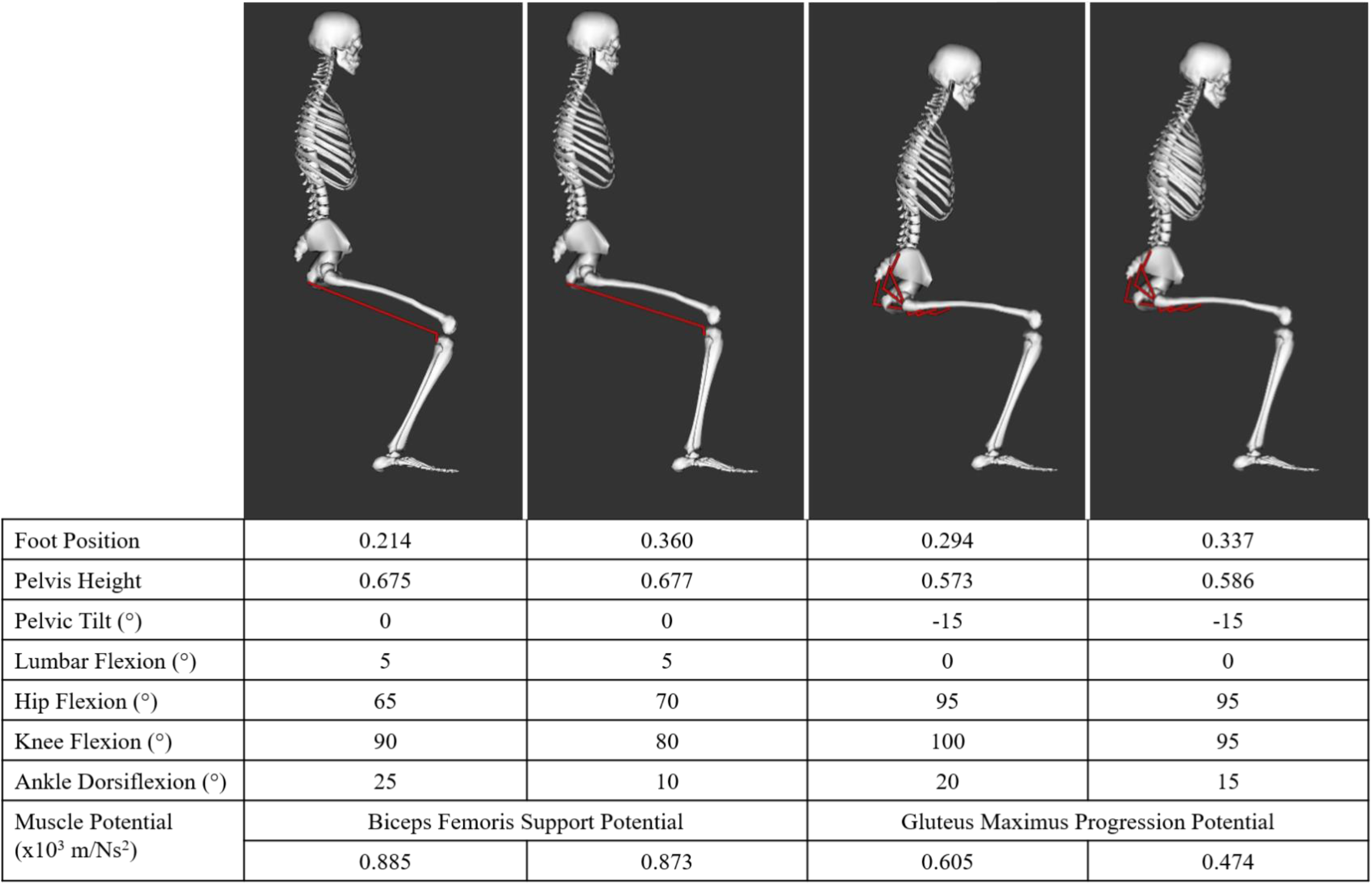
Select kinematic states and associated muscle potentials. Foot position and pelvis height were normalized to leg length.

The new method can be leveraged to explain actual kinematic adaptations used by individuals with KOA to complete the STS [21]. For example, individuals with KOA commonly demonstrate weakened quadriceps [22]. Therefore, it would be advantageous for these individuals to move in a way that increases the potential of the quadriceps to contribute to the STS while decreasing the force required to produce the necessary contribution to acceleration. With the greater anterior pelvic tilt observed in the individuals with KOA, their rectus femoris has greater potential for support during the momentum transfer phase when support demands increase significantly as the chair no longer supports the body.

Furthermore, quadriceps strengthening alone does not improve physical function in a considerable number of patients (up to 40%) [11]. With this new method it is also possible to explore alternative compensatory movement kinematics that may leverage other muscles to compensate for the weak muscle group and improve patient mobility. For example, severe knee pain in some individuals with KOA may inhibit treatments to strengthen the quadriceps or significantly alter the knee kinematics. These individuals could potentially improve their overall muscle contributions to support through kinematic modification by decreasing their anterior pelvic tilt angle by just 5° from −35° to −30°, to increase both gluteus maximus and biceps femoris support potentials by 50% and 60%, respectively. Additionally, for kinematic states with pelvic tilt angles of −30° to −20°, the gluteus maximus has, on average, up to 126.4% greater potential for support than rectus femoris, with greater differences between muscles with decreasing anterior pelvic tilt. Such an analysis could suggest that rehabilitation that combines movement training, to help a patient with KOA perform the STS with less than 30° of anterior pelvic tilt, with gluteus maximus muscle strengthening, to increase its peak isometric force, could increase muscle contributions to support.

An alternative method to modify STS kinematics is to increase the chair height [16], either by selecting a taller chair when available or using a cushion to boost the effective height. The chair height will be equal to the pelvis height at the start of the momentum transfer phase, which begins the moment the individual is no longer supported by the seat. Given the, on average, approximately 30° of anterior pelvic tilt and 15° of lumbar flexion of the individuals with KOA at the start of the momentum transfer phase, the combined effects of pelvis height and foot position on muscle support potentials can be determined (Figure 11). Raising the pelvis height while keeping the foot position constant increases the support potential of the gluteus maximus, vasti, gastrocnemius, and soleus. Although gastrocnemius and soleus support potentials increase most with a more posterior foot position, further increases in support potentials can be achieved by raising the pelvis height. Increasing pelvis height by 4.2% of leg length (3.3 cm for the generic 2D model) from 0.513 to 0.555 increases gastrocnemius and soleus support potentials by 10% and 11%, respectively. It is worth noting that increasing pelvis height would also increase the negative support potential of tibialis anterior. However, previous research has shown that tibialis anterior is not a primary contributor to the STS transfer [6,13], therefore its potential will not have a significant effect on contributions to support due to minimal tibialis anterior force production.

**Figure 11:**
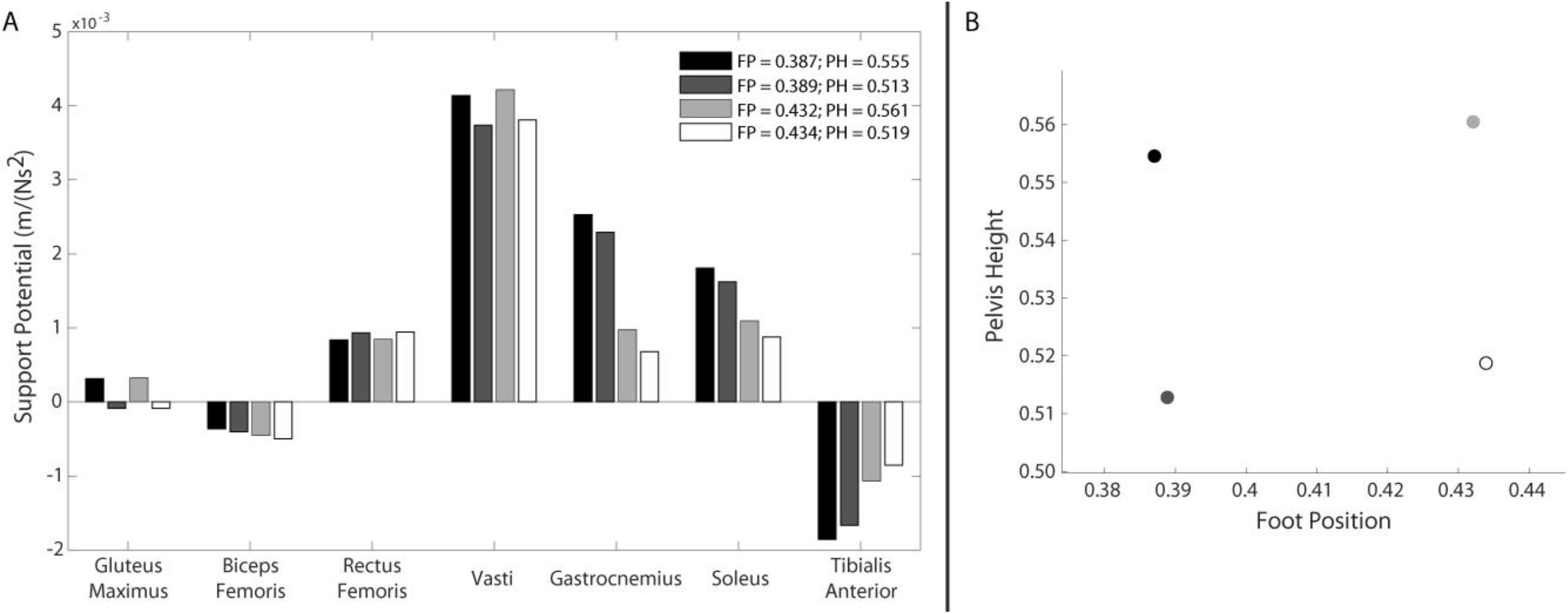
A) Influence of foot position and pelvis height on muscle support potentials for 30° of anterior pelvic tilt and 15° of lumbar flexion (average KOA kinematics at the start of the momentum transfer phase). B) Graphical depiction of selected pelvis heights and foot positions associated with muscle potentials in (A). Foot position and pelvis height were normalized to leg length.

In addition, rising from a higher chair would increase the quadriceps ability to perform their important functional role during the momentum transfer phase of resisting forward COM acceleration [6] and prevent a dangerous forward fall. Given the, on average, approximately 40° of anterior pelvic tilt and 10° of lumbar flexion of the individuals with KOA when the braking COM acceleration peaks in the second half of the momentum transfer phase, the new method determined that the quadriceps have the greatest potential to resist progression at greater pelvis heights and that a more anterior foot position further increases the vasti’s braking potential (Figure 12). Thus, for this example, the new method suggests that at a foot position of 0.300 and pelvis height of 0.706, the vasti have the greatest potential for braking and the plantarflexors, which are highly active during the momentum transfer phase due to their key role in providing support [6], have a smaller progression potential. Therefore, by increasing the potential of the vasti to contribute to braking and reducing the plantarflexor contribution to progression that the quadriceps would need to resist, this kinematic state would reduce the required quadriceps force, which are commonly weak in individuals with KOA. The analyses described could be used to identify appropriate seat height adjustments (e.g., adding a cushion to the seat to increase height) and initial foot position to reduce the required quadriceps load while providing sufficient support and resistance to a forward fall.

**Figure 12:**
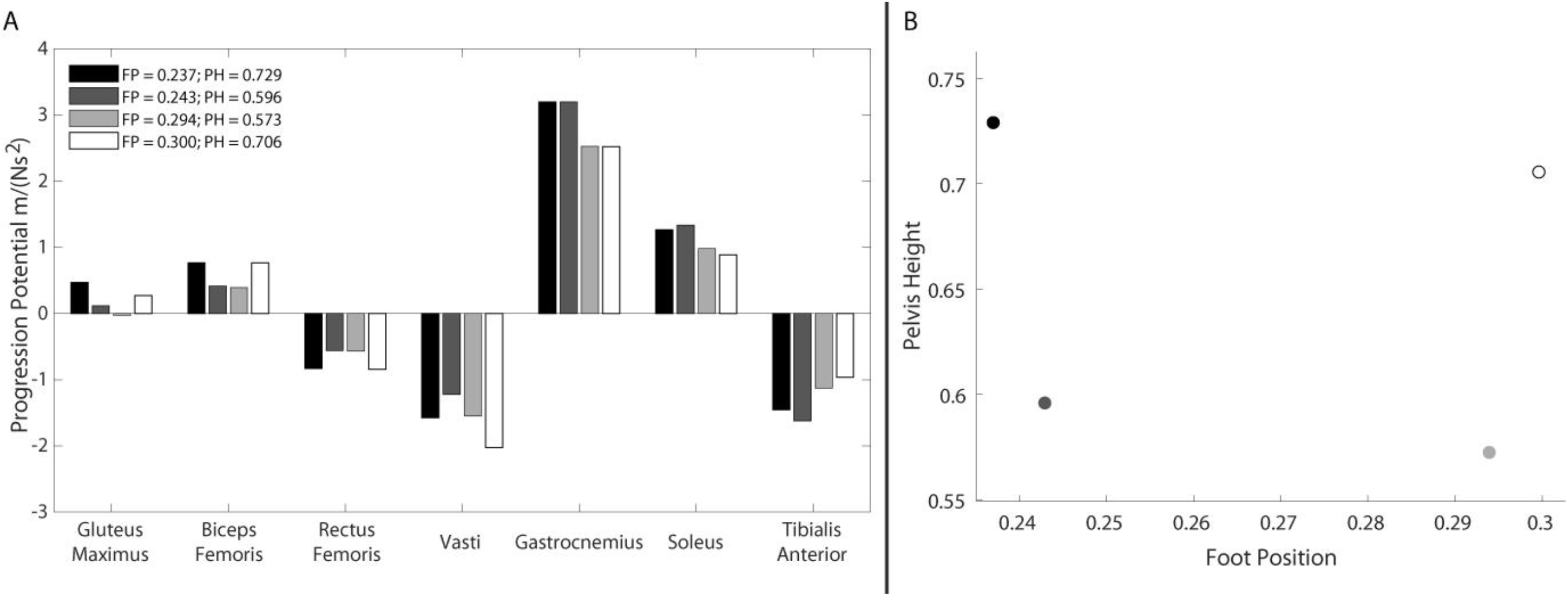
A) Influence of foot position and pelvis height on muscle progression potentials for 40° of anterior pelvic tilt and 10° of lumbar flexion (average KOA kinematics in the middle of the momentum transfer phase). B) Graphical depiction of selected pelvis heights and foot positions associated with muscle potentials in (A). Foot position and pelvis height were normalized to leg length.

This new method provides a foundation to be expanded upon to develop more extensive analyses. First, in its current formulation, the new method uses a static approach which does not account for joint velocity. However, the aim of this study was to systematically assess the effects of a range of different kinematic positions on muscle potentials, rather than retrospectively analyze a unique, experimentally measured movement with specific changes in joint positions over time. Future work is needed to adapt the current method to assess the effects of dynamic changes in joint positions on muscle potentials. In addition to the critical role of a muscle’s potential in determining its contribution to COM acceleration, the force produced by the muscle is an important factor determining a muscle’s contribution to the movement and also depends on joint kinematics. The location of a muscle on its force-length-velocity curve is determined by the angle of the joint(s) the muscle crosses and the change in joint position(s) over time. Although a muscle’s *potential* to contribute to acceleration is independent of the muscle’s force, the muscle’s contribution will depend on the muscle’s length, contraction velocity, and activation. Initial next steps will leverage this new method to determine the relative influence of joint kinematics on muscle potentials and the muscle force-length relationship. Such analyses could identify the optimal kinematic state for a muscle to produce the greatest contribution to COM acceleration for a given performance criterion (e.g., minimize force demand of a specific muscle or minimize muscle activation). To enhance opportunities for researchers to build on this initial work, the code for the new method is accessible on GitHub (https://github.com/sroelkerPhD/ipa-2d-4link).

## Conclusions

In conclusion, this work developed an efficient method to systematically evaluate the effects of sagittal plane kinematic modifications on individual muscle potentials, identify possible sources of pathologic movement, and identify potential rehabilitation strategies that leverage kinematics and muscle strength to improve mobility. We demonstrated how kinematic modifications at all joints affect a muscle’s ability to contribute to a movement through the role of joint positions in determining the mass matrix and muscle moment arms. The new method computes muscle potentials consistent with those estimated by simulation techniques while overcoming the time and resource limitations that impede the application of IAA in a clinical setting. When applied to the momentum transfer phase of the STS transfer, the new method revealed how the combined effects of kinematic modifications, including altering the pelvic tilt angle, adjusting the foot position, and increasing the chair height, could increase muscle potentials to leverage current muscle strength and also identified potential compensatory muscles that could be targeted for muscle strengthening in rehabilitation. Incorporating the principles of IAA into clinical practice has the potential to provide insight into how patients may benefit from both kinematic and muscle strength gains.

